# Uncovering a Sex-Specific cAMP Landscape in Vascular Smooth Muscle

**DOI:** 10.64898/2026.09.11.751051

**Authors:** Eric A Pereira da Silva, Ildernandes Vieira-Alves, Hannah Voorhees, Padmini Sirish, Nuria Daghbouche-Rubio, Miguel Martín-Aragón Baudel, Yang K. Xiang, Manuel F. Navedo, Madeline Nieves-Cintrón

**Author notes:** Correspondence: Madeline Nieves-Cintrón, Ph.D. Department of Pharmacology University of California, Davis, CA One Shields Avenue, Davis, CA 95616, Manuel F. Navedo, Ph.D. Department of Pharmacology University of California, Davis, CA, One Shields Avenue, Davis, CA 95616.

## Abstract

The second messenger, cyclic adenosine monophosphate (cAMP), plays a crucial role in regulating cellular function, including in vascular smooth muscle (VSM). While extensive research has focused on the compartmentalization of cAMP signaling in various cell types, the influence of biological sex on distinct cAMP pools and their functional implications in the VSM remains largely unexplored. To address this knowledge gap, we employed a multiscale experimental approach spanning targeted cAMP biosensors, wire myography, live-cell calcium imaging, proximity ligation assays, and *in vivo* ultrasound imaging. Our findings reveal a sex-specific distribution of cAMP signaling domains, with the cAMP pool selectively present in the SR of females but not males in the VSM. Ovariectomy abolished this sexual dimorphism, with SR cAMP pools disappearing in ovx females, mirroring the pattern observed in male cells. Males and ovx female VSM exhibited a significantly stronger RyR-PDE3/PDE4 association than sham females, suggesting that ovarian hormone-dependent sequestration of phosphodiesterases away from the SR licenses cAMP signaling in this compartment in female cells. Functionally, this sexually dimorphic cAMP compartmentalization correlates with smaller Ca^2+^ spark amplitude, reduced isoproterenol-induced relaxation, and higher pulse wave velocity in male and ovx compared to female VSM/vessels/mice. Together, these findings establish biological sex as a fundamental determinant of subcellular cAMP organization in VSM, with implications for understanding sex differences in vascular function and cardiovascular disease.

**GRAPHICAL ABSTRACT:** 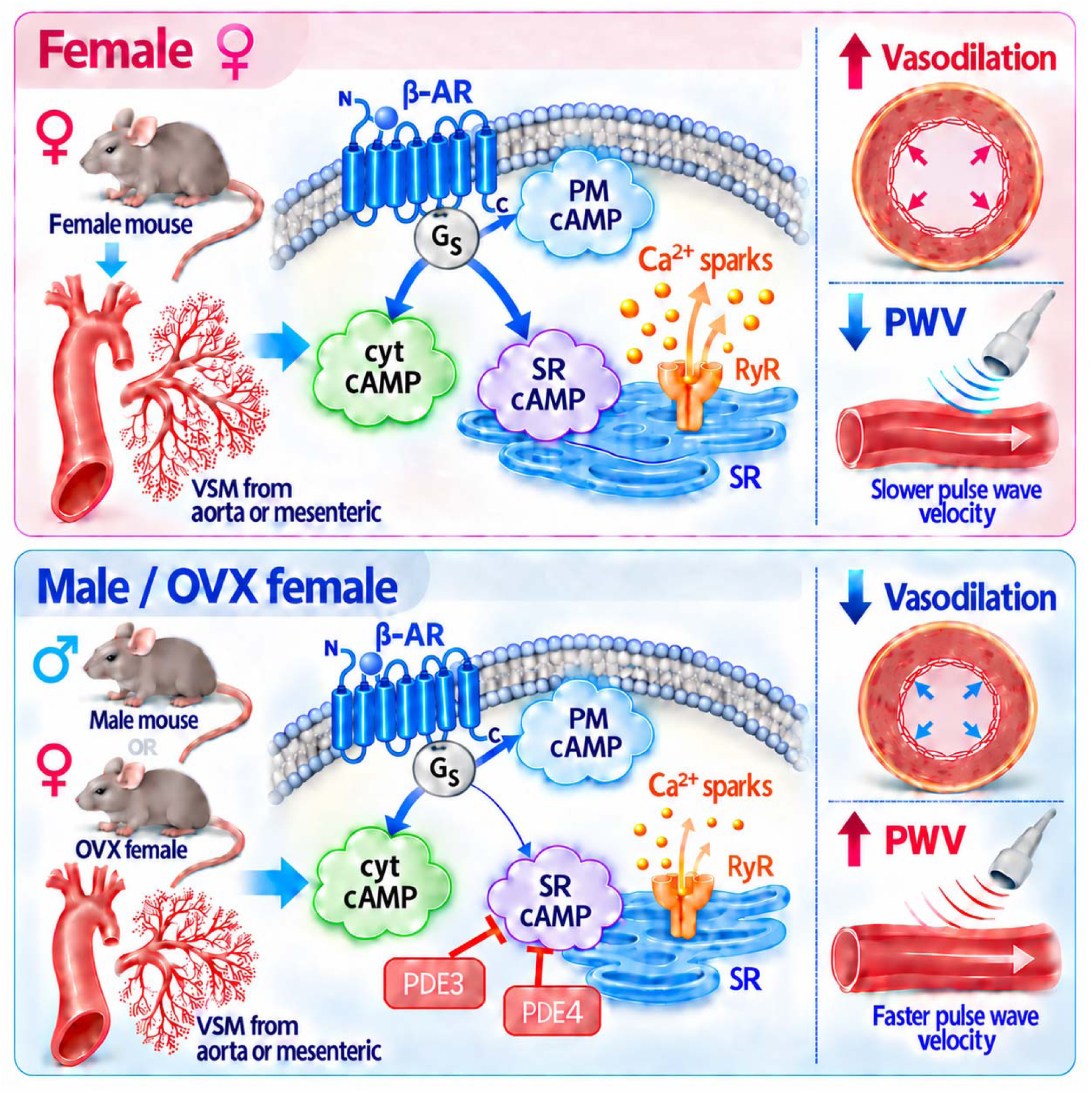

**Sex-dependent compartmentation of cAMP signaling in vascular smooth muscle.** In females, β-adrenergic stimulation generates robust plasma membrane, cytosolic, and sarcoplasmic reticulum-localized cAMP signals that support RyR-dependent Ca^2+^ sparks, vasodilation, and lower pulse wave velocity. In males and ovariectomized females, PDE3/PDE4 activity constraints sarcoplasmic reticulum cAMP pool, which correlates with reduced Ca^2+^ spark activity, reduced vasodilation, and increased pulse wave velocity.

**KEY POINTS:**

- Using compartment-targeted cAMP biosensors, we found that β-adrenergic stimulation raises cAMP at the plasma membrane and cytosol in vascular smooth muscle of both sexes but generates a sarcoplasmic reticulum cAMP pool only in female cells.
- Ovariectomy abolishes the female-specific pool, indicating that the sarcoplasmic reticulum response depends on ovarian hormones.
- Phosphodiesterase (PDE) transcript levels are similar across groups, but PDE3 and PDE4 sit closer to the ryanodine receptor in males and ovariectomized females, and inhibiting either PDE unmasks the sarcoplasmic reticulum cAMP pool.
- Absence of the sarcoplasmic reticulum cAMP pool in male and ovariectomized female VSM is accompanied by smaller Ca^2+^ spark amplitude, weaker isoproterenol-induced dilatation, and higher pulse wave velocity.
- Results suggest subcellular cAMP organization is a sex-dependent determinant of cellular function, vascular reactivity and arterial stiffness.

## INTRODUCTION

The ubiquitous second messenger 3’,5’-cyclic adenosine monophosphate (cAMP) is a critical regulator of cellular function, produced by adenylyl cyclase (AC) upon activation of stimulatory G protein-coupled receptors (G_s_PCRs). In vascular smooth muscle cells (VSM), cAMP serves as a central signaling node linking extracellular stimuli to vascular function, including excitation-contraction coupling (Hathaway et al., 1985; Orlov et al., 1996). For example, sympathetic activation and circulating vasodilators converge on G_s_PCRs such as β-adrenergic receptors (β-AR) to generate cAMP via AC stimulation, which drives protein kinase A (PKA) activation, membrane hyperpolarization via regulation of K^+^ channels, and ultimately relaxation of the vessel wall (Jackson, 2000; Nelson et al., 2011). Yet the view of cAMP as a uniformly relaxant signal in VSM has been challenged by work from our group demonstrating that engagement of a spatially distinct P2Y_11_-like/AC5 pathway triggers activation of a pool of PKA to promote L-type Ca_V_1.2 channel potentiation and VSM contraction in response to elevations in extracellular glucose (Hong et al., 2025; Prada et al., 2019; Syed et al., 2019). These findings revealed an unexpected contractile role for G_s_PCR-dependent cAMP signaling in the vasculature. Moreover, the results underscore a broader principle: cAMP’s functional output depends not solely on its abundance but on its precise subcellular localization. Nevertheless, whether VSM contains spatially distinct cAMP pools and what mechanisms govern their organization and functional outcomes remain unknown.

The versatility of cAMP as a second messenger is widely attributed to its spatial compartmentalization into discrete subcellular domains (Fu et al., 2024; Zaccolo and Kovanich, 2025). Evidence from other cell types, most notably cardiac myocytes, demonstrates that spatially confined cAMP pools are critical for regulating excitation-contraction coupling and organ function, raising the compelling possibility that analogous compartmentalization operates in VSM. Such compartmentalization is shaped by anchoring proteins, such as A-kinase anchoring proteins (AKAPs), that tether ACs and PKA, as well as by phosphodiesterase (PDE) enzymes, which hydrolyze cAMP and are key regulators of vascular reactivity (Fu et al., 2024) (Prada et al., 2020). In VSM, PDE3 and PDE4 are the predominant cAMP-hydrolyzing isoforms (Fu et al., 2024). Yet how PDEs’ subcellular localization shapes local cAMP pools in VSM remains unknown.

Compounding this gap in our understanding of compartmentalized cAMP is the poorly defined role of biological sex. Despite the overwhelming number of studies examining cAMP signaling in many cell types (Bers et al., 2019; Gold et al., 2013; Perera and Nikolaev, 2013; Sassone-Corsi, 2012; Singhrao et al., 2024), whether the spatial organization of discrete cAMP pools is sex-dependent has not been systematically investigated. Clues about sex-dependent changes in cAMP signaling come from experiments in heart cells and neurons. In cardiomyocytes, biochemical approaches revealed higher global cAMP levels in female than in male cells (Machuki et al., 2019). This was correlated with reduced PDE activity, resulting in increased L-type channel Ca_V_1.2 activity in female cells. In neurons, sex-specific differences in corticotropin-releasing factor receptor signaling and trafficking were associated with robust cAMP signaling in females but not males, and this was suggested to increase female vulnerability to stress-related psychopathology (Bangasser et al., 2010). Hormonal fluctuations in premenopausal women have been associated with differences in vasodilatory responses to cAMP-inducing agents such as β-AR agonists compared to men, and the differences are attenuated after menopause (Barnes, 2017; Freedman et al., 1987; Hart et al., 2011). Yet, underlying mechanisms, particularly in VSM, remain unclear.

Here, we tested the central hypothesis that biological sex shapes subcellular cAMP compartmentalization in VSM and that this organization contributes to vascular function. To address this, we employed a multiscale experimental approach combining targeted cAMP biosensors with live-cell FRET imaging, proximity ligation assays, live-cell calcium imaging, wire myography, and *in vivo* ultrasound imaging in VSM and vessels isolated from male, intact female, and ovariectomized (ovx) female mice. Our findings reveal sexually dimorphic cAMP compartmentalization that is ovarian hormone-dependent, mechanistically linked to differential PDE3 and PDE4 distribution at the sarcoplasmic reticulum (SR), and functionally consequential for calcium spark amplitude, vasodilation, and arterial stiffness. Together, results establish biological sex as a determinant of subcellular cAMP organization and vascular function.

## METHODS

### Animals

All animal procedures were conducted in full accordance with protocols approved by the Institutional Animal Care and Use Committee at the University of California, Davis. Eight-week-old male and female C57BL/6J mice were obtained from the Jackson Laboratory and group-housed in a 12:12-hour light-dark cycle with controlled humidity and temperature. Animals had ad libitum access to standard chow and water. Mice were euthanized by intraperitoneal injection of sodium pentobarbital (250 mg/kg), consistent with the approved University of California, Davis Institutional Animal Care and Use Committee protocol.

### Ovariectomy

Ovariectomy (ovx) was performed in female mice at 8-10 weeks of age. Mice were anesthetized with 2-5% isoflurane delivered by inhalation. Bilateral obx was performed through a single dorsal midline skin incision. The skin was gently separated from the underlying muscle to expose the lateral abdominal wall, and the ovaries were identified by visualizing the associated fat pads on each flank. Small muscle incisions, approximately 0.5 cm, were made over each fat pad, and the fat pads were gently exteriorized using fine forceps. The ovaries were clamped and excised, and the ovarian vessels were ligated at the uterine horn using 4-0 Vicryl sutures. Hemostasis was confirmed before the uterine horns were returned to the abdominal cavity. The muscle layers were closed using 3M Vetbond tissue adhesive (cat. no. 1469SB), and the skin was closed using 9-mm AutoClip clips (Fine Science Tools, cat. no. 12020-00). Mice received ketoprofen at 5 mg/kg preoperatively and once daily for two days postoperatively. After surgery, animals recovered in a clean cage with supplemental heat and were monitored until fully ambulatory before being returned to the housing room. Skin clips were removed 7 days after surgery. Five weeks after ovx, mice were euthanized for experimental analyses. Blood was collected, and uterine-to-body weight ratios were measured in sham-operated and ovx females to confirm surgical efficacy.

### Cytology

Vaginal smears from sham females and ovx mice were obtained by gently washing the vaginal opening 5x with 25 μl of PBS using a pipette. Immediately after, 5 μl of the fluid was added to a glass microscope slide. The smears were allowed to air-dry completely. Next, the slides were stained with 0.1% crystal violet (w/v) in distilled water, then submerged for 1 min in a Petri dish containing distilled water (2X). Once completely dried, a droplet of 100% glycerol was added to the slides, and a coverslip was placed. Slides were analyzed immediately using an inverted Micromaster microscope (Fisher Scientific) with infinity-corrected optics under bright-field (transmitted-light) illumination, equipped with 10× and 20× phase-contrast objectives and an integrated digital camera. Images were acquired using the manufacturer’s imaging software.

### Hormone Analysis

The steroid assay was performed by the West Coast Metabolomics Center at UC Davis. Blood samples were collected and placed into a chilled EDTA tube. To separate the plasma, the blood was centrifuged for 1 min at 3000 rcf, followed by 5 min at 10,000 rcf at 4°C. The plasma was stored at -70°C until analysis. Steroids from plasma were extracted in an antioxidant solution (0.2 mg/mL of butylated hydroxytoluene (BHT)/EDTA solution in 1:1 (methanol: water) by homogenizing in a GenoGrinder for 2 × 30 seconds, centrifuging, followed by washes in appropriate solvents, and lyophilizing. Samples were reconstituted for liquid chromatography-mass spectrometry (LC-MS) in 100 µL of 1 µM 1-phenyl-3-hexadecanoic acid urea/1-cyclohexyluriedo-3-dodecanoic acid (PHAU/CUDA) in methanol/acetonitrile (50:50), mixed, and then sonicated for 5 min, followed by centrifugation for 5 min in spin filters. The supernatant was transferred to glass inserts in a high-performance liquid chromatography HPLC vial. Tritiated steroids were added to monitor retention time, and steroids were recovered via HPLC, while deuterated internal standards were added to enable quantification of the compound of interest. To calculate the quantity of the steroid metabolite in each fraction, the area under the sample peak was divided by the area under the deuterated internal standard peak, and the result was reported as ng/mL for plasma levels. For estradiol analysis, blood samples were collected and processed as described above. Estradiol concentrations were determined in 50 µl plasma using a Mouse Estradiol Rapid ELISA Kit (Invitrogen, cat. no. EELR013), according to the manufacturer’s instructions. All samples were analyzed in duplicate. Estradiol levels were normalized to protein content using Pierce™ BCA Protein Assay (Thermo Fisher Scientific) and expressed as pg/μg.

### Aortic VSM Isolation

Primary, unpassaged vascular smooth muscle (VSM) cell cultures were established from isolated mouse aortas. Glass coverslips (#0; Karl Hecht, Sondheim, Germany) were coated with a 1:100 dilution of laminin (Fisher Scientific, cat. no. 23017015) prepared in sterile, calcium-free Dulbecco’s phosphate-buffered saline (DPBS; Gibco, cat. no. 14190144). The laminin-coated coverslips were placed in 24-well plates and incubated at 37°C in 5% CO_₂_ for at least 2 h. Aortas were dissected and transferred to dish with ice-cold dissection medium consisting of Dulbecco’s Modified Eagle Medium supplemented with GlutaMAX, 1 g/L D-glucose, and 110 mg/L sodium pyruvate (DMEM; Gibco Life Technologies, cat. no. 10567-014), as well as 1% penicillin-streptomycin (Gibco, cat. no. 15140) and 0.25% Fungizone (Invitrogen, cat. no. 15290018). Excess perivascular adipose and connective tissues were carefully removed. The aortas were then transferred to a Petri dish containing DMEM supplemented with 2.2 mg/mL collagenase type II (Worthington, cat. no. LS004174) and incubated at 37°C for 10 min to facilitate removal of the adventitia. Following incubation, the adventitial layer was removed using fine forceps. The aortas were opened longitudinally, and the luminal surfaces were gently swabbed with a cotton applicator to remove the endothelium. The denuded aortas were cut into small pieces and transferred to a 15-mL bioreaction tube with a vented cap (Neta Scientific, cat. no. CT229472). The tissue was incubated for approximately 2 h at 37°C with constant agitation in digestion buffer containing, in mM, 134 NaCl, 6 KCl, 1 MgCl_₂_, 2 CaCl_₂_, 10 HEPES, and 7 D-glucose, supplemented with 2.2 mg/mL collagenase type II.

Digestion was terminated by adding an equal volume of cultivation medium consisting of DMEM supplemented with 10% fetal bovine serum (FBS; Sigma, cat. no. F0926) and 1% penicillin-streptomycin. The digested tissue was centrifuged at 0.3 × 1000 rcf for 5 min, and the supernatant was discarded. The tissue pellet was resuspended in 60 µl of cultivation medium and gently mixed to disperse individual VSM cells. Before cell seeding, the laminin-coated coverslips were washed three times with sterile-filtered DPBS. 60 µl of the cell suspension was carefully applied to each coverslip, and the cells were incubated at 37°C in 5% CO_₂_ for 2 h to promote attachment. An additional 500 μL of cultivation medium was then added to each well. The cultures were maintained for 5-7 days, or until confluent, before viral infection.

### Adenovirus Transduction

Following the incubation period of 5-7 days and after cells reached ∼80% confluence, the media were replaced with 500 μL of serum-free media containing the virus encoding an Epac1-camps-based FRET sensor (ICUE3) or the cAMP Universal Tag for imaging experiments (CUTie), targeted to the plasma membrane (PM), cytosol (cyt), or sarcoplasmic reticulum (SR). Cells were incubated for 36-48 hours to allow infection, after which the medium was replaced with serum- and virus-free medium. The glass coverslips were transferred to glass-bottom culture dishes (MatTek) containing 3 mL of phosphate-buffered saline (PBS) at RT for experiments.

### Live-cell FRET Imaging of Isolated VSM

Live-cell FRET images were captured using an Olympus IX81 inverted fluorescence microscope (EVIDENT, Waltham, MA) equipped with a Photometrics Prime 95 cMOS camera (Tucson, AZ) and controlled by Metafluor software v7.7 (Molecular Devices). FRET images were acquired using a 60X water-immersion objective. For FRET analysis, the donor fluorophore was excited at 430-455 nm, and emission fluorescence was measured using two filters: 475DF40 for cyan and 535DF25 for yellow. Background subtraction was applied to the images, acquired every 20 s with a 200 ms exposure time per channel. The donor/acceptor FRET ratio was calculated and normalized to the baseline ratio in the absence of stimulation. Results are expressed as the average change in FRET ratio induced by exposure to 100 nM ISO, relative to the maximal response elicited by 10 μM forskolin (FSK) + 100 μM 1-Methyl-3-Isobutylxanthine (IBMX). All experiments were conducted at room temperature (22-25°C).

### Flow cytometry analysis

VSM cell suspensions were passed through a 200-µm cell strainer and fixed with 0.4% paraformaldehyde. Cells were then incubated overnight at 4°C with Alexa Fluor 488-conjugated anti-α-smooth muscle actin (Abcam, Cambridge, MA), phycoerythrin-conjugated anti-Thy1.2, anti-CD31, and anti-CD45 antibodies, and a lineage antibody cocktail containing antibodies against CD3ε, CD11b, CD45R, Ly-6C, Ly-6G, and TER-119 (BD Biosciences). Antibody staining was performed in PBS containing 5% serum and 20 µg/mL DNase-free RNase (Sigma). Cells were also stained with 40 µg mL^−1^ 7 aminoactinomycin D (7AAD, BD Biosciences, San Jose, CA) to assess DNA content. Data were collected using a BD LSRII flow cytometer. Data were analyzed using BD FlowJo 10.10.0 software.

### Quantitative Polymerase Chain Reaction

Mesenteric arteries were isolated, lysed, and RNA extracted using SPLIT RNA Extraction Kit (Lexogen; cat. No. K00848). RNA was reverse transcribed with High-Capacity RNA-to-cDNA Kit (Applied Biosystems; cat. no. 4387406). RT product was used for quantitative PCR. PDE transcript expression was analyzed in mesenteric arterial lysates using Power SYBR Green PCR Master Mix (Applied Biosystems; cat. no. 4367659). Specific primers were used to detect PDE2A (*Pde2a*; NM_001243758.2), PDE3A (*Pde3a*; NM_018779), PDE3B (*Pde3b*; NM_011055), PDE4A (*Pde4a*; NM_019798), PDE4B (*Pde4b*; NM_019840), PDE4D (*Pde4d*; NM_011056). The PDE primers were synthesized by Integrated DNA Technologies (IDT) using the previously reported primer sequences (Carvalho et al., 2021). A commercially available primer assay for GAPDH (*Gapdh*; NM_008084; cat. no. 249900) was acquired from Qiagen (Valencia, CA). Amplification was performed using an Applied Biosystems real-time PCR instrument. Expression for each gene was normalized to GAPDH.

### Proximity Ligation Assay (PLA)

Duolink in situ proximity ligation assay (PLA) was used to examine the colocalization between RyR2, PDE3, and PDE4 proteins in freshly isolated mesenteric and aortic arterial myocytes from male, female sham, and ovx. Dissociated myocytes were plated on glass coverslips and allowed to adhere for an hour at room temperature (RT). Arterial myocytes were washed once with PBS, fixed in 3% glyoxal solution (Sigma-Aldrich, cat # 128465) for 20 min, quenched in 100 mM glycine (Sigma-Aldrich, cat # 50046) in PBS for 15 min, then washed 3X for 5 min with PBS. Cells were permeabilized using 0.1% Triton X-100 for 20 min and subsequently blocked in 5% Intercept blocking buffer/0.1% Triton X-100 blocking solution (LI-COR Biosciences, cat # 927-70001) for 1 hour at RT. Cells were incubated overnight with the following primary antibodies combinations: mouse anti-RyR2 (Invitrogen #MA3-916, 1:200), rabbit anti-PDE3A (Abcam #ab99236, 1:200), rabbit anti-PDE4 (Abcam #ab14628, 1:200). These antibodies were diluted (1:200) in a solution containing 0.05% Triton X-100, 0.01% Intercept blocking buffer in PBS, mixed with Duolink Antibody Diluent (cat. no. DUO82008) at 1:1 ratio. Cells were incubated with only one primary antibody as a negative control. Following overnight incubation with primary antibodies at 4 °C, cells were washed 3X times for 5 min with Buffer A. PLA rabbit PLUS and mouse MINUS probes were used to detect different protein combinations (1:5 in Duolink Antibody Diluent, 1 hour at 37 °C). Cells were washed with Duolink Buffer A 3X for 5 min, then incubated with the ligation solution for 30 min at 37°C to allow hybridization and formation of a circular DNA template at sites of dual labeling. This was followed by 3 washes of 5 min each in Buffer A, and cells were incubated in amplification solution for 100 min at 37°C, followed by 2X 10 min washes with Duolink Buffer B and 1X 1 min wash in high-purity H_2_O. Finally, coverslips were allowed to dry, then mounted on a microscope slide with Duolink mounting medium containing DAPI (cat. no. DUO82040) and sealed with coverslip sealant (Biotium, cat. no. 23005).

For imaging, we used an Andor Revolution spinning-disk confocal system coupled to an Olympus iX-81 inverted microscope equipped with an Olympus 60X oil-immersion objective (NA 1.4), an Andor iXon EMCCD Ultra camera, 561- and 405-nm lasers, and appropriate excitation and emission filters. The system was controlled using Andor IQ 2.8 software. Confocal z-stacks were acquired at 0.5 μm steps while maintaining a constant acquisition rate, exposure time, number of z-steps, and laser power across experimental conditions. Maximum-intensity projection images generated from the combined z-stacks were used to analyze the number of puncta per cell volume (μm^3^). Image analysis was performed using Imaris x64 v10.1.1 (Oxford Instruments) by team members blinded to experimental conditions.

### Calcium (Ca^2+^) sparks recording

Vascular smooth muscle cells (VSM) were obtained from mesenteric arteries of male, sham-operated female, and ovx female mice. Mesenteric arteries were dissected in ice-cold dissection buffer containing (in mM): 140 NaCl, 5 KCl, 2 MgCl2, 10 D-glucose, and 10 HEPES, pH 7.4 adjusted with NaOH. After dissection, arteries were incubated in dissection buffer supplemented with papain (1 mg/mL) and dithiothreitol (1 mg/mL) at 37 °C for 9.5 min. Arteries were then incubated in a dissection buffer containing collagenase type H (1.77 mg/mL), elastase (0.5 mg/mL), and trypsin inhibitor (1 mg/mL) for 9.5 min at 37 °C. Following enzymatic digestion, arteries were washed 3X in ice-cold dissection buffer, followed by 2 washes in an ice-cold dissociation buffer containing (in mM): 125 NaCl, 5.4 KCl, 15.4 NaHCO3, 0.33 Na2HPO4, 0.44 KH2PO4, 3 Sucrose, 10 D-Glucose, and 11 HEPES, pH 7.4 adjusted with NaOH and supplemented with 100 nM CaCl_2_. After the last wash in dissociation buffer, the tissue was gently triturated with polished glass pipettes to promote the dissociation of individual VSMs. Isolated VSMs were maintained in ice-cold solution until imaging.

VSMs were placed on a circular 25-mm #1.5 coverslip in a recording chamber for 10 min to allow adhesion. The cells were loaded with the Ca^2+^-Sensitive fluorescent indicator Cal 520-AM (AAT Bioquest, cat. no. 3031542). Cal-520 AM stock solution was prepared fresh in DMSO and diluted 1:200 in recording solution to a final concentration of 5 μM. VSMs were incubated in the dye for 5-10 min to allow indicator loading, then washed with recording solutions containing (in mM): 140 NaCl, 5 KCl, 1 MgCl2, 2 CaCl2, 10 D-Glucose, 10 HEPES, pH 7.4 with NaOH. Recordings were performed at room temperature. An Andor spinning-disk confocal microscopy system coupled to an Olympus iX81 inverted microscope equipped with a 60× oil-immersion objective was used to image Ca^2+^ sparks. Images were acquired at 90-110 Hz with an 8 ms exposure using the Andor IQ software. Spark movies were analyzed using SparkAn software, developed by A. D. Bonev and M. T. Nelson (University of Vermont). Thirty images without spark activity were averaged to determine baseline fluorescence levels (F_0_). Regions of interest of 7 pixels^2^, corresponding to approximately 4 μm^2^, were used to detect sparks using a minimum F/F_0_ threshold of 1.2 and a 20% tolerance. Large Ca^2+^ events, including waves, were excluded from the analysis. When the same spark event appeared across multiple frames, only one event was retained for analysis. Distinct spark events within the same cell were averaged for analysis.

### Wire myography

Wire performed wire myography experiments using a DMT Multi Wire Myograph System (model 620), connected to an isometric force transducer. Transducer signals were digitized using an 8-channel PowerLab analog-to-digital converter and recorded using LabChart software (AD Instruments, Hastings, UK). Myograph baths were filled with modified Krebs buffer (PSS in mM): 119 NaCl, 4.7 KCl, 1.2 KH_2_P0_4_, 1.2 MgCl_2_, 24 NaHC0_3_, 0.03 EDTA, 7 Glucose, and 2 CaCl_2_. The solution was maintained at 37°C and continuously aerated with 95% O_2_ and 5% CO_2_.

Second- and third-order mesenteric artery segments were dissected in ice-cold MgPSS buffer (in mM): 140 NaCl, 5 KCl, 2 MgCl_2_, 10 glucose, and 10 HEPES, pH 7.4. Arteries were allowed to stabilize for 15 min, then subjected to a passive force normalization procedure. Arteries were constricted with 60 mM KCl-PSS, followed by a wash to allow tension to return to baseline. Acetylcholine (ACh; 10 μM; Sigma-Aldrich, cat. no. A2661) was used to assess endothelial function. Arteries were considered endothelium-denuded when ACh-induced relaxation was <20%. Arteries were precontracted with phenylephrine (Phe; 2 μM; Sigma-Aldrich, cat. no. 1001166140). Arteries were precontracted with 1 μM Phe, and cumulative concentration-response curves were generated using (-)-isoproterenol hydrochloride (ISO; Sigma-Aldrich, cat. no. I6504), with concentrations ranging from 0.001 nmol/L to 10 μM. Between ISO concentrations, arteries were allowed to equilibrate for 5 min before applying the next concentration.

For aortic experiments, the thoracic aorta was carefully excised and placed in ice-cold Krebs buffer. Connective tissue was carefully removed, and the aorta was cut into 3-4 mm segments. Aortic segments were gradually stretched to a resting tension of 10 mN over 10 min and allowed to equilibrate for 10 min. Vessels were initially contracted with 60 mM KCl-Krebs solution for 5 min, then washed with Krebs solution to allow tension to return to baseline. This procedure was repeated with a 10-min contraction in 60 mM KCl-Krebs solution, followed by 2 additional washes with Krebs. Acetylcholine (ACh; 10 μM; Sigma-Aldrich, cat. no. A2661) was used to assess endothelial function. Arteries were considered endothelium-denuded when ACh-induced relaxation was <20%.Arteries were precontracted with 1 μM Phe, and cumulative concentration-response curves were generated using (-)-isoproterenol hydrochloride (ISO; Sigma-Aldrich, cat. no. I6504), with concentrations ranging from 0.001 nM to 10 μM. Between ISO concentrations, arteries were allowed to equilibrate for 5 min before applying the next concentration.

For analysis, we selected 120-second segments from LabChart and exported absolute tension values to Microsoft Excel. Data were then converted to percentage responses, normalized to Phe contraction.

### Pulse wave velocity (PWV)

Pulse wave velocity (PWV) was measured using the Vevo F2 High-Resolution *In Vivo* Imaging System (VisualSonics, Toronto, ON, Canada) to assess aortic stiffness. Data were analyzed using Vevo LAB software v5.11.1. Briefly, anesthesia was induced by placing the mouse in an induction chamber with 2-5% isoflurane and 100% oxygen at 2 L/min for 2 min. Once the animal lost its righting reflex, it was placed supine on a heated platform, with its nose inserted into a nose cone to maintain anesthesia with 1.5% isoflurane. The limbs were secured to the platform ECG electrodes using electrode cream to monitor heart rate, ECG, and respiratory rate throughout imaging. Hair over the anterior chest and abdomen was removed using depilatory cream, and warmed ultrasound gel was applied to the imaging area. PWV was measured from B-mode and pulsed-wave Doppler images of the abdominal aorta. Pulsed-wave Doppler recordings were acquired sequentially from proximal and distal locations along the abdominal aorta. Color Doppler imaging was used as needed to confirm blood flow direction and optimize Doppler beam alignment.

During offline analysis, the time interval from the onset of the ECG QRS complex to the foot of the Doppler velocity waveform was measured for both the proximal and distal recording sites. Mice with heart rate > 400 bm were used for analysis. The distance between the two Doppler sampling locations was measured along the centerline of the abdominal aorta on the corresponding B-mode image. Pulse transit time was calculated as the difference between the distal and proximal QRS-to-foot intervals, and PWV was calculated by dividing the distance between the two sampling sites by the pulse transit time. Analyses were performed using the built-in PWV module in Vevo LAB software.

### Statistics

Data are presented as mean ± SEM unless otherwise indicated. Statistical analyses were performed using GraphPad Prism. Data distribution was assessed using a normality test before statistical analysis. For comparisons between two groups, normally distributed data were analyzed using an unpaired Student’s *t* test, whereas non-normally distributed data were analyzed using the Mann-Whitney test. For comparisons among more than two groups, normally distributed data were analyzed using one-way ANOVA followed by an appropriate post hoc multiple-comparisons test. Non-normally distributed data involving multiple groups were analyzed using the Kruskal-Wallis test followed by Dunn’s multiple-comparisons test. *P* < 0.05 was considered statistically significant.

### Chemicals

All chemical reagents were from Sigma-Aldrich (St. Louis, MO) unless otherwise stated.

## RESULTS

### Sexually dimorphic cAMP signaling in VSM

To examine cAMP signaling domains in VSM, we used live-cell FRET imaging using cAMP biosensors (ICUE3 and CUTie; Figure 1A and 1B, respectively) targeted to the cytosol (cyt), plasma membrane (PM), and sarcoplasmic reticulum (SR) (DiPilato et al., 2004; DiPilato and Zhang, 2009; Reddy et al., 2018; Surdo et al., 2017). The ICUE3 and CUTie biosensors report elevated cAMP via decreases or increases in the FRET ratio, respectively (DiPilato et al., 2004; DiPilato and Zhang, 2009; Reddy et al., 2018; Surdo et al., 2017). The subcellular localization of each targeted biosensor has been previously validated (Reddy et al., 2022). We expressed the sensors in unpassaged primary aortic VSM (Figure 1C). This approach was used given the challenges of expressing exogenous proteins in native vascular cells and the current lack of mouse models expressing compartment-specific cAMP biosensors. While primary culture inevitably involves some degree of phenotypic modulation from contractile to synthetic states, our established protocol preserves key features of native VSM, including elongated cell morphology, robust high potassium-induced elevation in intracellular calcium, and high VSM purity as confirmed by flow cytometry showing >98% positivity for α-smooth muscle actin and absence of endothelial, fibroblast, and other lineage markers (Figure 1C, 1D) (House et al., 2008; Prada et al., 2020; Syed et al., 2019).

**Figure 1:**
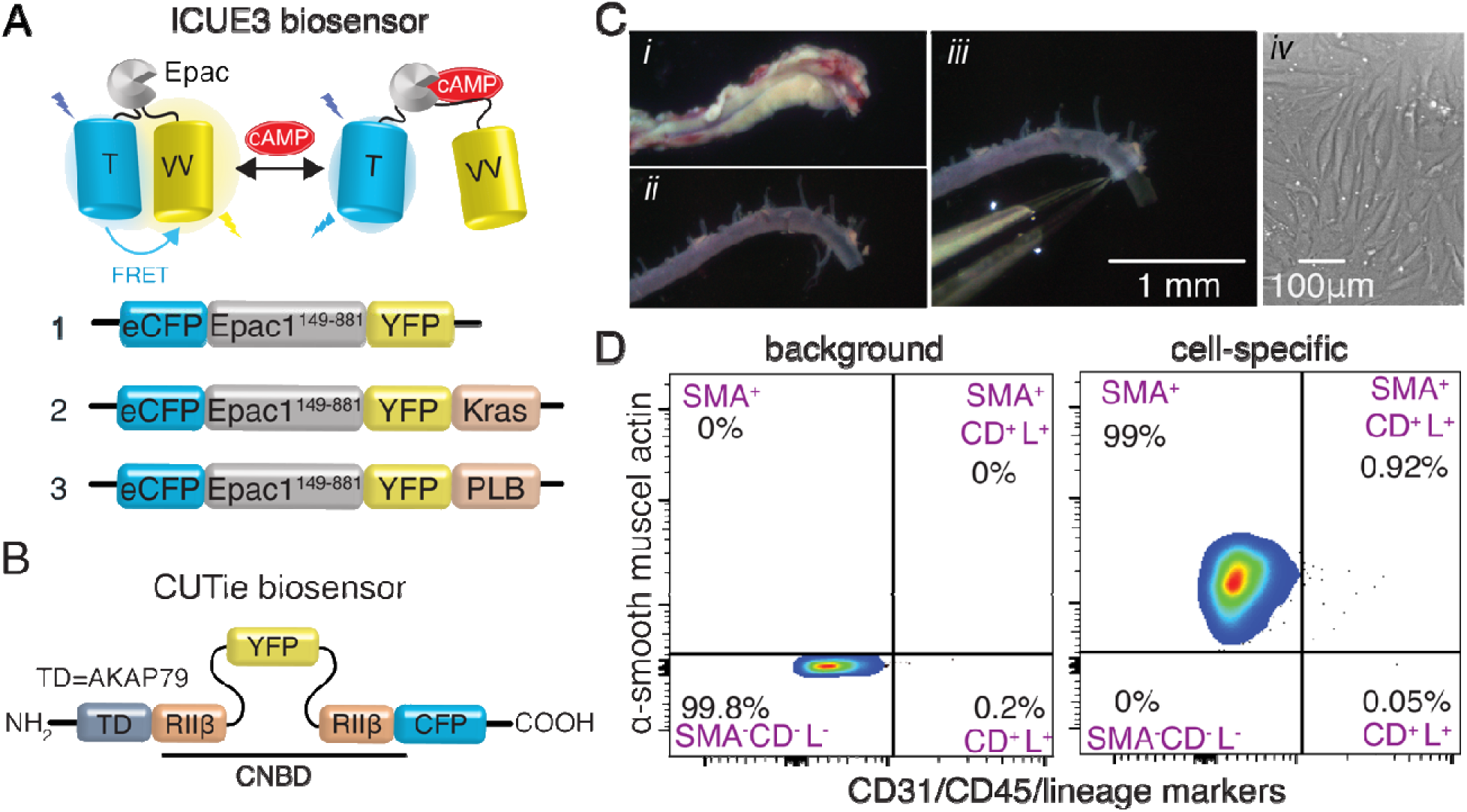
Isolation and characterization of primary mouse aortic smooth muscle cells. **A)** Cartoon of ICUE and **B)** CUTie cAMP biosensors. **C)** Workflow for establishing unpassaged VSM cultures; (*Ci*) aorta freshly dissected from a mouse, (*Cii*) aorta after removal of the surrounding fat tissue, (*Ciii*) removal of the adventitia from the medial layer after enzyme incubation, C*iv)* representative image of cultured aortic smooth muscle cells (10X objective). **D)** Flow cytometric analyses of smooth muscle cells: (left) background fluorescence with isotype controls, (right) Smooth muscle cells identified as α-smooth muscle actin-positive, CD31-, CD45-, lineage-negative (α-SMA+/Thy-/CD31-/CD45-/Lin-). X- and Y-axes represent arbitrary units. Representative results are shown. N = 9 mice.

To examine cAMP pools, the β-AR receptor was used as a prototypical G_S_PCR to stimulate cAMP production in VSM. For this, β-AR was stimulated with the β-AR agonist isoproterenol (ISO, 100 nM). In male VSM, application of ISO decreased the FRET ratio in the PM and cytosolic compartments, reflecting a rise in local cAMP (Figures 2A, 2B), but produced no detectable response in the SR (Figure 2C). Subsequent application of the broad adenylyl cyclase activator forskolin (fsk) combined with the pan-PDE inhibitor IBMX confirmed sensor functionality by elevating cAMP in all compartments. These data indicate that β-AR stimulation generates spatially restricted cAMP signals preferentially at the PM and in the cytosol, while the SR compartment remains insulated from this response in male VSM.

**Figure 2:**
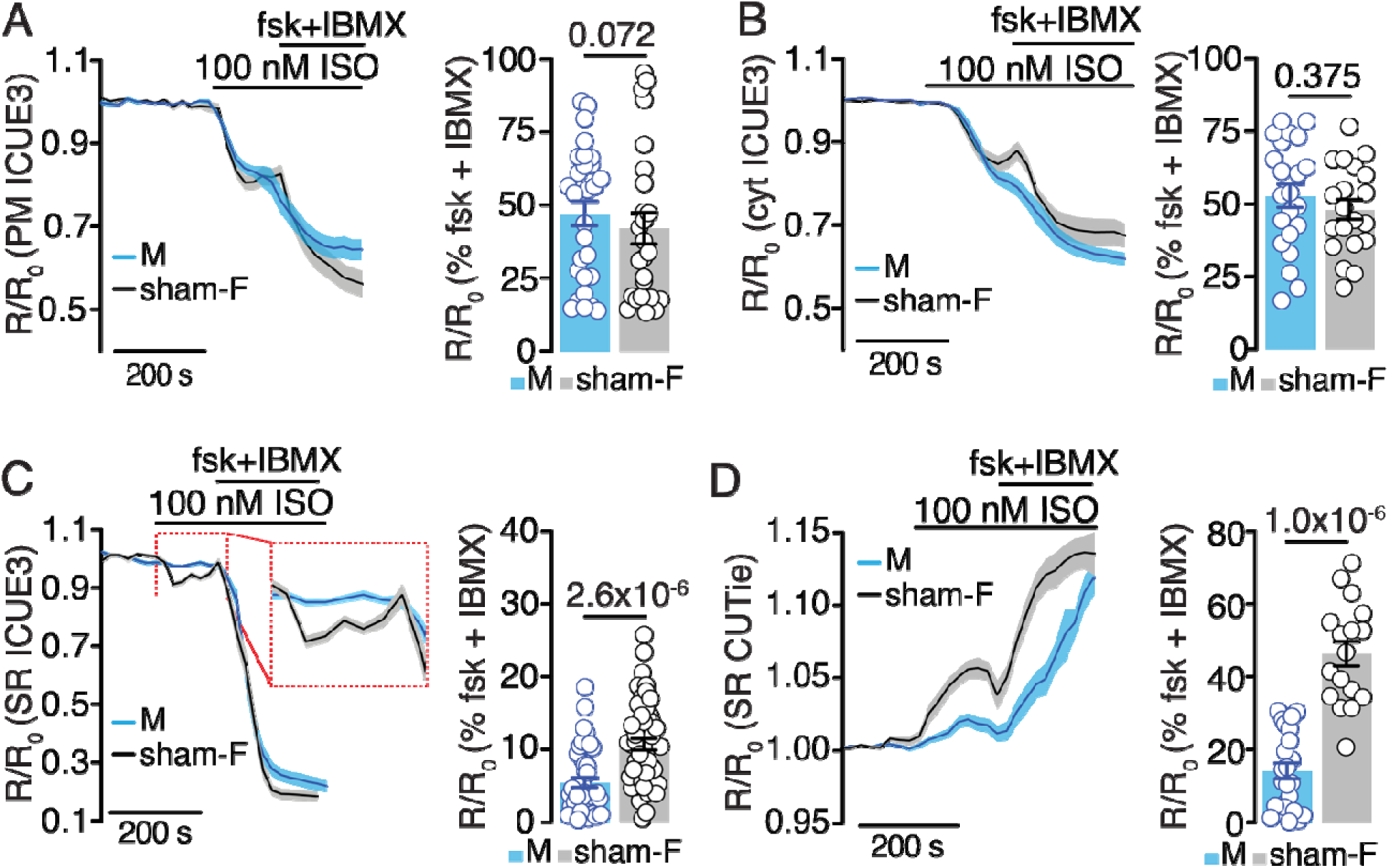
Sexually dimorphic cAMP signaling domains in VSM. (n=cells/arteries; N= mice) **A-C)** Time courses of normalized FRET ratios in response to isoproterenol (ISO), followed by the maximal response to forskolin plus the phosphodiesterase inhibitor IBMX, and summary plots of ISO-induced FRET responses expressed as a percentage of the forskolin + IBMX response. cAMP responses were measured using targeted ICUE3 sensors at the **A)** plasma membrane (PM), **B)** cytosol (Cyt), and **C)** sarcoplasmic reticulum (SR) in VSMs from male and sham-operated female mice. **D)** Time courses and summary plots of SR-localized cAMP responses measured using the CUTie sensor in VSMs from male and sham-operated female mice. Data are presented as the mean ± SEM. Sample sizes are as follows (n=cells and N= cultures): ICUE3 experiments: male: PM (n/N= 29/5); Cyt (n/N= 22/8); SR (n/N 45/3); sham: PM (n/N = 22/7), Cyt (n/N= 21/5), SR (n/N= 51/4). CUTie experiments, male (n/N= 25/6); sham ( n/N = 17/10). *P*-values are shown in the figure. Statistical analysis by Kruskal-Wallis test followed by Dunn’s multiple-comparisons test.

To determine whether this compartmentalization differs between sexes, we compared ISO-evoked responses in male versus sham-operated female VSM. At the PM and cytosol, ISO produced comparable FRET responses in male and sham female VSM, with no significant difference between groups (Figures 2A, 2B). However, the SR compartment revealed a pronounced sex difference, with ISO eliciting a robust decrease in the FRET ratio in sham female VSM compared to male VSM (Figure 2C). Consistent with this finding, imaging with the orthogonal SR-targeted sensor CUTie confirmed that ISO-evoked SR cAMP signals were significantly larger in female compared to male VSM (Figure 2D). These results suggest that β-AR-stimulated cAMP accumulation at the SR is greater in female VSM. Together, these results reveal a sex-specific difference in SR cAMP compartmentalization, whereby female VSM exhibits preferential β-AR-driven cAMP accumulation in the SR, which is absent or markedly reduced in males.

### Ovarian hormones are required for sexually dimorphic SR cAMP signaling in VSM

Next, we tested the role of ovarian hormones in establishing the sexually dimorphic SR cAMP compartments. Age-matched female littermates were randomized into a sham surgery group (surgery without ovary removal; sham) or an ovariectomized group (ovx) (Figure 3A). Five weeks post-surgery, hormonal status was confirmed by vaginal cytology, and the uterine-to-body weight (BW) ratio was calculated as an additional index of ovary removal and ovarian hormone depletion. As expected, ovx displayed vaginal cytology predominantly composed of leucocytes, with a significant reduction in the uterine-to-BW ratio compared with the sham-F group (Figures 3B and 3C). Circulating levels of estradiol, progesterone, and its metabolite, 20 α-dihydroprogesterone (20 α-DHP), were significantly lower in ovx than in the sham group (Figure 3D-3F). These results confirmed successful ovarian hormone depletion.

**Figure 3:**
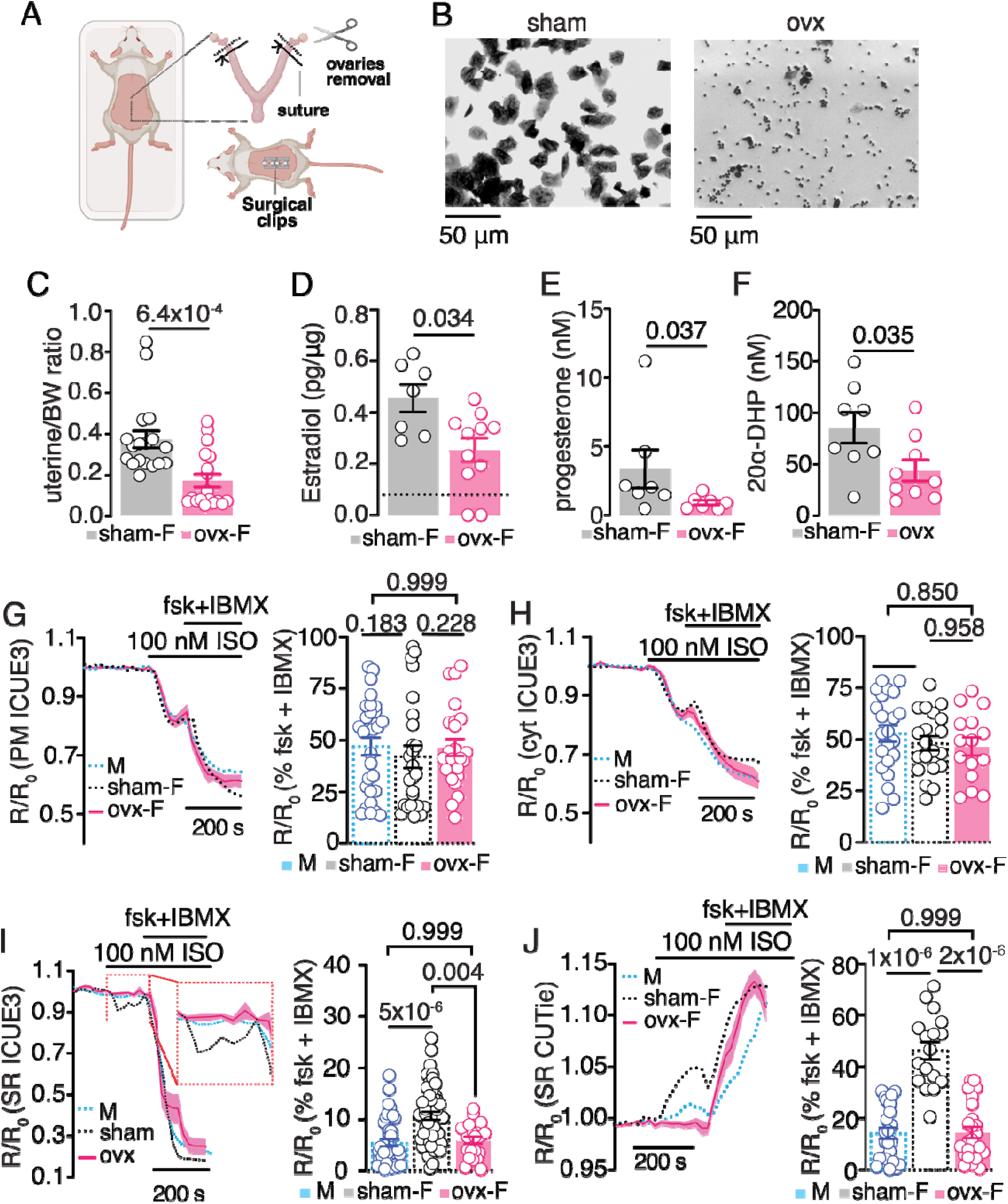
Role of ovarian hormone loss on compartmentalized cAMP signaling. **A)** Cartoon of the ovariectomy procedure. **B)** Representative brightfield images of vaginal cytology stained with 0.1% crystal violet from (i) a sham-operated mouse, showing nucleated and cornified epithelial cells, and (ii) an ovariectomized (ovx) mouse, showing predominantly leukocytes. Images were acquired using a 20× objective. **C-F)** Effects of ovariectomy on the **C)** uterine/body weight ratio (sham N=18; ovx N=18); and estradiol (sham N= 7; ovx N=11); **D)** progesterone (sham N=7; ovx N=7); and **E)** 20-dihydroprogesterone (20-DHP) (sham N=8; ovx N=9) **(F)** levels (N=mice). Data are presented as the mean ± SEM. *P*-values are shown in the figure; Statistical analysis: Mann-Whitney. **G-J)** Time courses of normalized FRET ratios in response to isoproterenol (ISO), followed by the maximal response to forskolin plus the phosphodiesterase inhibitor IBMX, and summary plots of ISO-induced FRET responses expressed as a percentage of the forskolin + IBMX response. cAMP responses were measured at the **G)** plasma membrane (PM), **H)** cytosol (Cyt), and **I, J)** sarcoplasmic reticulum (SR) in VSMs from male, sham-operated female, and ovx mice. The male, sham, and ovx experiments were performed as part of the same experimental series, with recordings generally acquired in parallel under matched experimental conditions. Male and sham-operated female data are also presented in Figure 2 and are reproduced here as dotted traces to permit direct comparison with the ovx group; these data were included in the statistical analyses shown in this figure. Sample sizes are as follows (n=cells and N= cultures): Plasma membrane: male (n/N= 29/5); sham (n/N= 22/7); ovx (n/N 23/3). Cytosol: male (n/N= 22/8); sham (n/N= 21/5); ovx (n/N= 18/3). SR_ICUE3: male (n/N= 45/3); sham (n/N= 51/4); ovx (n/N= 22/3). SR_CUTie: male (n/N= 25/6); sham (n/N= 17/10); ovx (n/N= 25/9). Data are presented as the mean ± SEM. *P*-values shown in the figure. Statistical analysis: PM, SR-ICU3 and SR-Cutie; Kruskal-Wallis test followed by Dunn’s multiple-comparisons test; Cyt; one-way ANOVA, followed by Tukey’s multiple comparison test.

Primary VSM cultures were then established from ovx mice and transduced with compartment-targeted biosensors. Male and sham female VSM data from Figure 2 are replotted as dotted lines alongside the ovx data to facilitate direct comparison. In response to ISO application, ovx cells displayed a significant increase in cAMP at the PM and in the cytosolic compartment, comparable to that observed in male and sham VSM (Figure 3G, 3H). Strikingly, the cAMP response in the SR compartment of ovx cells was markedly reduced, mirroring the pattern observed in male rather than sham female VSM (Figure 3I, 3J). Together, these findings demonstrate that the preferential β-AR-driven cAMP accumulation at the SR observed in female VSM requires intact ovarian hormone signaling and that its loss recapitulates the male phenotype.

### Role of PDEs on SR cAMP signaling in VSM

PDEs are key regulators of cAMP spatiotemporal dynamics (Maurice et al., 2003; Zaccolo, 2011; Zhang et al., 2019). We first asked whether differential PDE abundance could account for sex-dependent SR cAMP responses. Transcript analysis confirmed expression of PDE2A, PDE3, PDE4A, PDE4B, and PDE4D in vascular tissue from males, sham females, and ovx females, with no significant differences among groups (Figure 4A). These data indicate that global PDE transcript expression does not explain the divergent SR cAMP responses. Instead, we hypothesized that differences in the subcellular organization of PDE activity may explain the differences. While direct SR-anchored PDE complexes are well characterized in cardiac muscle, their organization in VSM is unclear (Fu et al., 2024).

**Figure 4:**
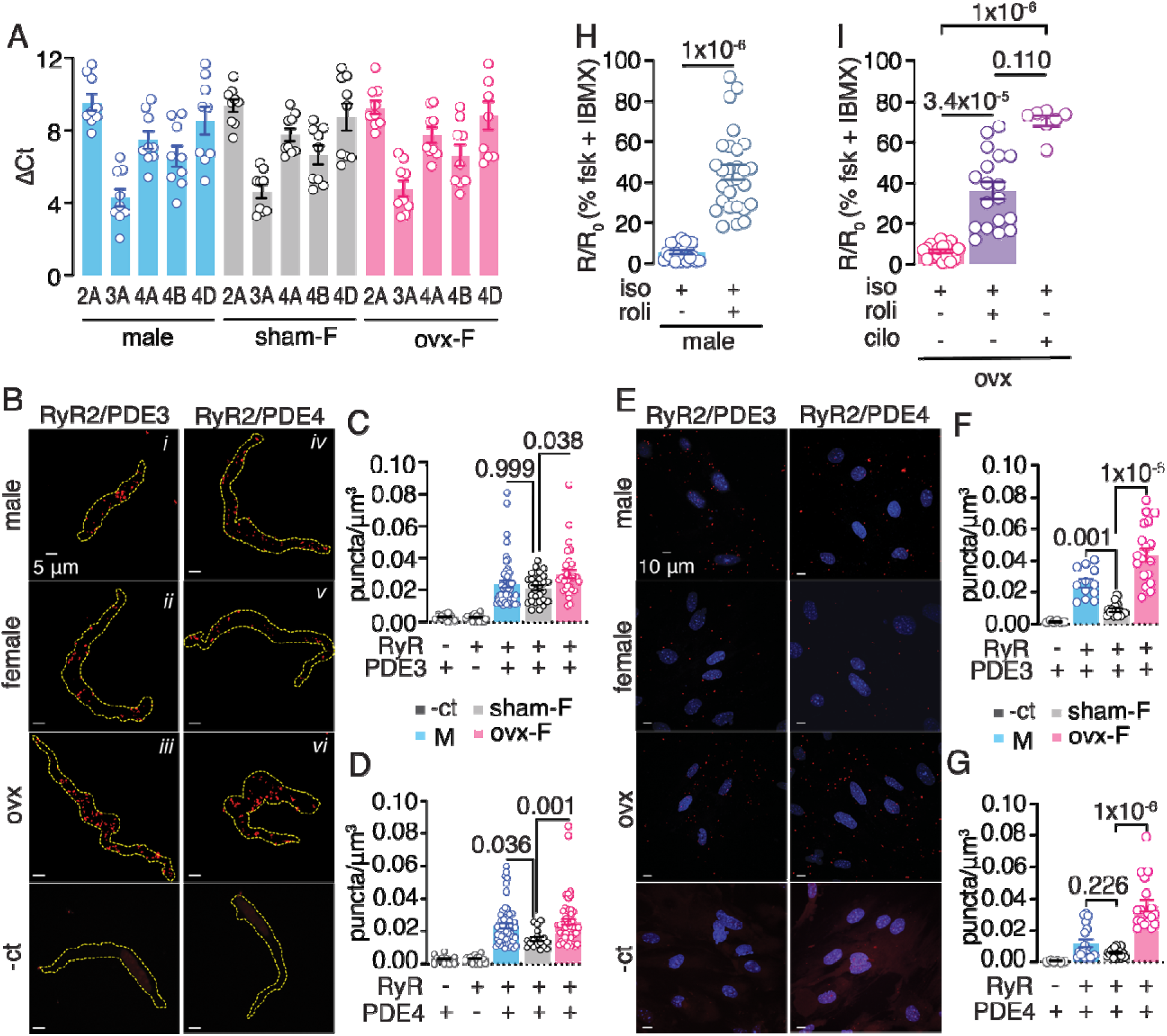
Association of PDE3 and PDE4 with the SR of VSM. **A)** Relative expression of PDE isoforms in mesenteric arterial lysates from male (N=9), sham (N=9), and ovx (N=9) mice, presented as ΔCT values. **B)** Representative fluorescence images and quantification of proximity ligation assay (PLA) puncta/μm^3^ in acutely isolated mesenteric VSM co-labeled for **C)** RyR2 and PDE3 or **D)** RyR2 and PDE4. Representative images are shown for male (i, iv), sham-operated female (ii, v), and ovx female (iii, vi) mice. Sample size (n=cells and N= mice). RyR2-PDE3 PLA: male (n/N= 59/7); sham (n/N 28/6); ovx (n/N = 31/6). For RyR2-PDE4 PLA: male (n/N= 56/6); sham (n/N= 21/5); ovx(n/N= 57/7). **E)** Representative fluorescence images and quantification of PLA puncta per μm^3^ in unpassaged aortic vascular smooth muscle cells co-labeled for **F)** RyR2 and PDE3 or **G)** RyR2 and PDE4. Sample size: RyR2-PDE3 PLA: male (n/N=12/3); sham (n/N= 15/4); ovx ( n/N= 18/3). RyR2-PDE4 PLA: male (n/N= 18/5); sham (n/N= 19/4); ovx (n/N=18/5). Data represent mean ± SEM; *P-values* are shown in the figure; one-way ANOVA with Šidák’s multiple comparisons. **H)** The PDE4 inhibitor rolipram unmasks sarcoplasmic reticulum-localized cAMP responses in male VSM and restores these responses in ovx VSM. Sample sizes (n=cells and N= cultures). Male: - roli (n/N= 45/3); +roli (n/N= 27/3). Data are presented as the mean ± SEM. *P*-values shown in the figure. Statistical determined statistical significance using a t-test. **I)** The PDE3 inhibitor cilostamide restores SR-localized cAMP responses in ovx VSM. ovx: +ISO (n/N= 22/3); ISO+roli (n/N= 18/2); ISO+cilo (n/N= 7/2). Data are presented as the mean ± SEM. *P*-values shown in the figure; Kruskal-Wallis test.

PDE3 and PDE4, constitute the predominant cAMP-hydrolyzing activities in VSM (Hubert et al., 2014; Lugnier et al., 1986; Maurice et al., 2003; Singhrao et al., 2024), are the principal isoforms coupled to β-AR cAMP signaling (Zhai et al., 2012) and have been shown to anchor to SR macromolecular complexes (PDE3A within SERCA2-AKAP18δ-PKA complexes; PDE4D within mAKAP-RyR complexes) (Ahmad et al., 2015; Lehnart et al., 2005). Therefore, we focused on these PDEs to determine their potential contributions to the observed phenotype. To determine whether PDE3 and PDE4 isoforms are physically close to the SR in VSM, we employed the proximity ligation assay (PLA), which detects proteins within 40 nm of each other (Fredriksson et al., 2002). For these experiments, we immunostained VSM with antibodies against the SR marker ryanodine receptor (RyR) and either PDE3 or PDE4 to assess their proximity to the SR membrane. Negative controls using a single primary antibody produced no detectable PLA signal, confirming assay specificity in both acutely isolated and unpassaged cultured cells. In acutely isolated VSM from mesenteric arteries, PLA revealed RyR-PDE3 and RyR-PDE4 associations in both males and sham females (Figures 4B-4D). Importantly, RyR-PDE4 proximity signals were significantly greater in male than in sham female VSM (Figure 4C, 4D). In ovx females, both RyR-PDE3 and RyR-PDE4 puncta were significantly elevated compared to sham females (Figures 4B-D), indicating that ovarian hormone depletion shifts the PDE4 landscape at the SR toward the male pattern and increases PDE3 association. Complementary findings were obtained in unpassaged primary-cultured aortic VSM, where RyR-PDE3 and RyR-PDE4 PLA signals were detected in both sexes (Figure 4E-G). RyR-PDE4 proximity signals were trended higher in male than in sham female VSM, but the difference was not significant. Consistent with the acutely isolated data, both RyR-PDE3 and RyR-PDE4 puncta were significantly higher in ovx than in sham female VSM (Figures 4E-G).

To directly test the functional consequence of SR-localized PDE activity, we tested whether PDE inhibition could restore the blunted ISO-evoked SR cAMP response observed in males and ovx females VSM expressing the SR ICUE3 sensor (Figure 4H, 4I). The selective PDE4 inhibitor rolipram (10 nM) significantly increased ISO-induced SR cAMP accumulation in both male and ovx female VSM, consistent with the greater PDE4 association with the SR detected by PLA. Moreover, in ovx female VSM, the PDE3-selective inhibitor cilostamide also rescued the SR cAMP response (Figure 4I), consistent with increased PDE3 localization to the SR in this group. Taken together, these findings reveal a sex hormone-dependent mechanism of SR cAMP compartmentalization in VSM, whereby ovarian hormones suppress PDE4 association with the SR, thereby permitting robust β-adrenergic-stimulated cAMP accumulation in this domain in female VSM. Loss of ovarian hormones, whether naturally absent as in males or experimentally induced by ovariectomy, shifts the PDE landscape toward greater SR-localized PDE3 and PDE4, effectively insulating the SR cAMP pool from β-AR stimulation.

### Role of distinct PDE distribution on Ca^2+^ sparks in VSM

RyR-mediated Ca^2+^ release events (Ca^2+^ sparks) are key regulators of VSM contractility whose activity is tuned by local cAMP/PKA signaling (Jaggar et al., 1998; Tykocki et al., 2017). Although sex differences in Ca^2+^ entry and global intracellular Ca^2+^ have been reported in VSM, direct Ca^2+^ spark-level comparisons between sexes remain unclear (Asuncion-Alvarez et al., 2024; O’Dwyer et al., 2020). Given that PDE3 and PDE4 are physically associated with RyR at the SR, we reasoned that their differential distribution across groups could locally restrict cAMP availability and thereby modulate Ca^2+^ spark activity. To test this, we recorded Ca^2+^ sparks in freshly isolated mesenteric arterial myocytes loaded with the Ca^2+^ indicator Cal 520-AM (Figures 5A-F).

**Figure 5:**
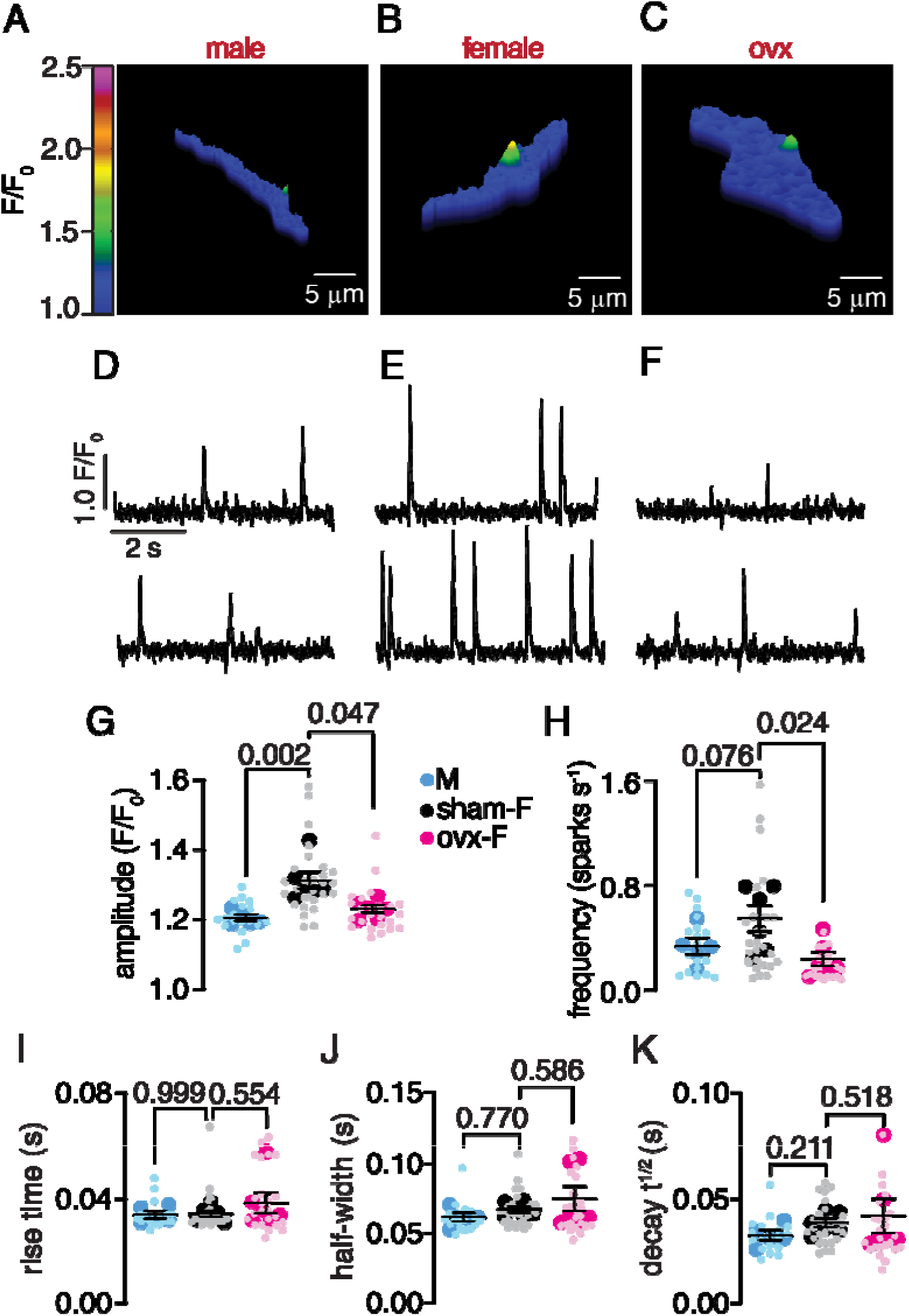
Distinct SR PDE distribution is associated with altered RyR-mediated Ca^2+^ signaling in VSM. **A-C)** Representative pseudocolor images of VSM showing Ca^2+^ spark events and **D-F)** representative spark recording traces from male, sham female, and ovx mice. Summary plots show **G)** Ca^2+^ spark amplitude, **H)** frequency**, I)** rise time, **J)** half-width, and **K)** decay time in cells from male (N = 5), sham female (N = 6), and ovx (N = 6) mice. Data are presented as the mean ± SEM for each animal, with measurements for individual cells (small dots) overlaid. *P*-values are shown in the figure. Statistical tests were performed using the Kruskal-Wallis test, followed by Dunn’s multiple-comparisons test.

Ca^2+^ spark amplitude was significantly higher in VSM from sham females than in males and females (Figure 5G). Ca^2+^ spark frequency did not differ between males and sham females but was significantly reduced in ovx females compared with both groups (Figure 5H). Other kinetic parameters did not differ significantly among groups, including rise time, half-width, and decay time (t_1/2_) (Figures 5I-K). Together, these results establish a functional correlation between compartmentalized PDE activity, SR cAMP pools, and RyR-mediated Ca^2+^ signaling in VSM and identify a novel hormonally regulated mechanism by which ovarian hormones shape the Ca^2+^ spark landscape in VSM.

### Males and ovx females exhibit reduced β-AR-induced vasodilation compared to sham females

Having established that SR cAMP signaling is selectively enhanced in sham females and dependent on ovarian hormones, we next asked whether this difference translates into functionally distinct vasodilatory responses to β-AR stimulation. To test this, we assessed ISO-induced relaxation in mesenteric arteries from males, sham females, and ovx females using wire myography (Figure 6). Mesenteric arteries were mounted in a wire myography chamber and stimulated with 60 mM KCl to assess basal contractility, followed by precontraction with phenylephrine (Phe, 2 µM). We then applied cumulative doses of ISO to assess β-AR-induced dilation of precontracted vessels. KCl-induced contraction did not differ between groups (Figure 6A). While Phe-induced contractions were similar between males and sham females, ovx females exhibited significantly higher contractile responses (Figure 6B). ISO-induced dilation of mesenteric arteries was higher in sham females than in males or ovx females (Figure 6C). To determine whether this pattern extends to a conduit vessel, we examined ISO-induced relaxation in thoracic aortic rings from the same three groups. KCl-induced contraction was significantly greater in sham females than in males or ovx females, while Phe-induced (1 µM) contractions were similar across groups (Figure 6D and E, respectively). Mirroring the mesenteric findings, aortic rings from sham females displayed significantly greater dilation in response to ISO than those from males or ovx females (Figure 6F). Together, these results demonstrate that ovarian hormones are required for enhanced β-AR-induced vasodilation in both resistance and conduit vessels, a functional pattern that mirrors the sex- and hormone-dependent differences in SR cAMP compartmentalization observed at the cellular level.

**Figure 6:**
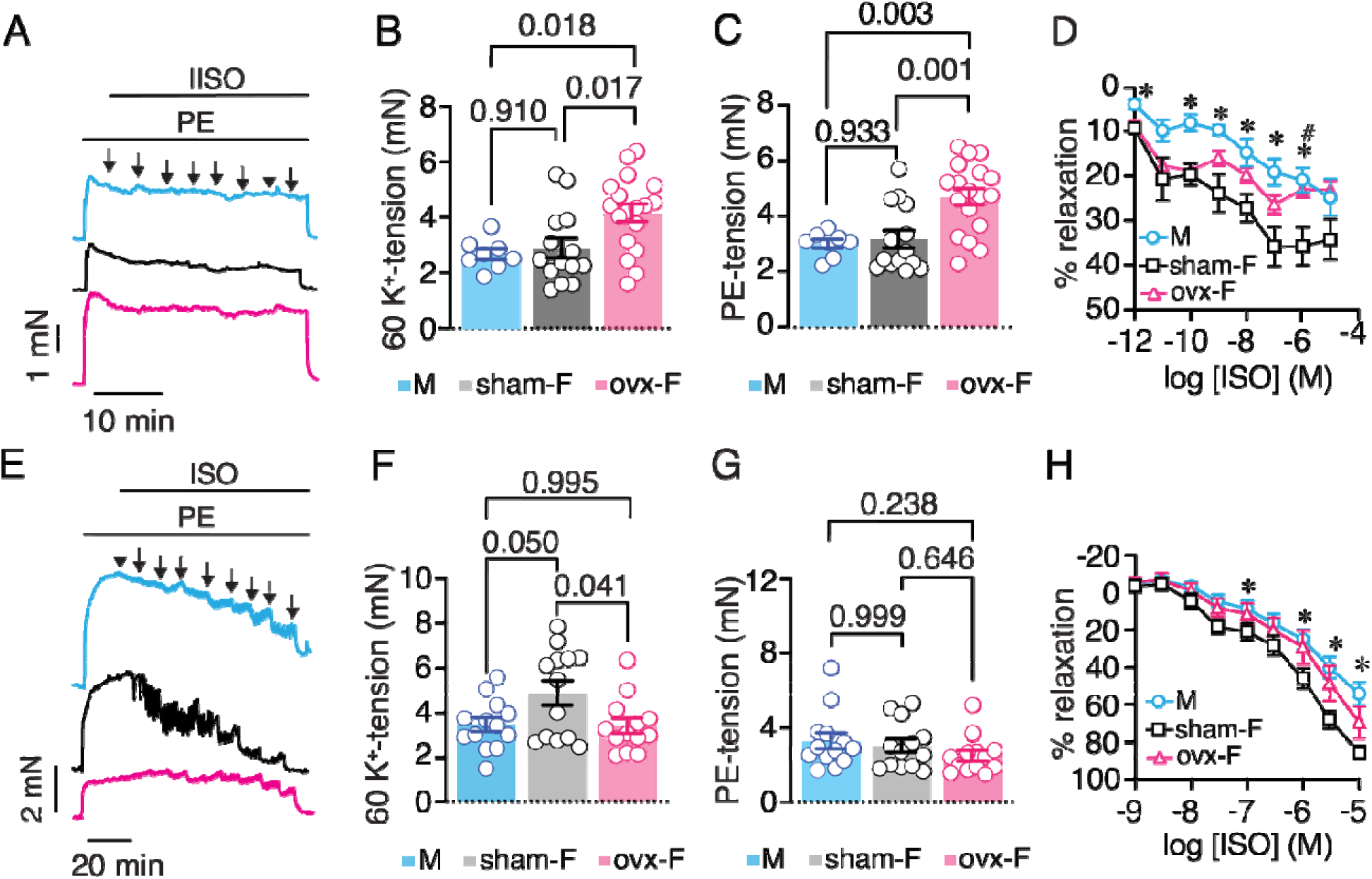
Sexually dimorphic vasomotor response in the mesenteric arteries and thoracic aorta. **A)** Representative myography traces recorded in mesenteric arteries from male (top), sham female (middle), and ovx female (bottom). Summary plots of **B)** 60mM KCl-induced tension and **C)** Phe-induced tension in mesenteric arteries from male, sham female, and ovx mice. **D)** Concentration-response curve for ISO-induced dilation of mesenteric arteries precontracted with 2 μM Phe. Responses were normalized to the Phe-induced precontraction and are expressed as percent relaxation. Sample sizes: male 8; sham 13; ovx 23. Data are presented as the mean ± SEM. *P values* are indicated in the figure; one-way ANOVA with Šidák’s multiple comparisons test. For panel D, \**P* < 0.05, male versus sham; and #*P* < 0.05, sham versus ovx; Mixed-effects analysis followed by Dunnett’s multiple-comparisons test. **E)** Representative myography traces recorded in thoracic aorta rings from male (top), sham female (middle), and ovx female (bottom). Summary plots of **F)** 60 mM KCl- induced tension and **G)** Phe-induced tension in aortic rings from male, sham-operated female, and ovx mice. **H)** Concentration-response curves for ISO-induced dilation of aortic rings precontracted with 1 μM Phe. ISO concentrations ranged from 1 nM to 10 μM. Responses were normalized to the Phe-induced preconstriction and are expressed as percent dilation. Sample sizes: male, 13; sham, 17; and ovx, 13 mice. Data are presented as the mean ± SEM. *P* < 0.05, male versus sham; one-way ANOVA followed by Tukey’s multiple-comparisons test.

### Sex and ovarian hormones influence arterial stiffness

Sex and ovarian hormones influence vascular compliance and hemodynamics (Novella et al., 2012; Ogola et al., 2018). Thus, we assessed pulse wave velocity (PWV) and aortic Doppler parameters in male, sham female, and ovx female mice to determine whether the cellular and tissue differences described above are accompanied by changes in arterial stiffness. Representative ultrasound and Doppler recordings used for pulse transit-time measurements are shown in Figures 7Ai and 7Aii. PWV was significantly higher in ovx females than in sham females, indicating increased arterial stiffness following ovarian hormone depletion (Figure 7B). Heart rate did not differ significantly among groups (Figures 7C). Together, these data indicate that loss of ovarian hormones promotes a vascular phenotype characterized by increased arterial stiffness.

**Figure 7:**
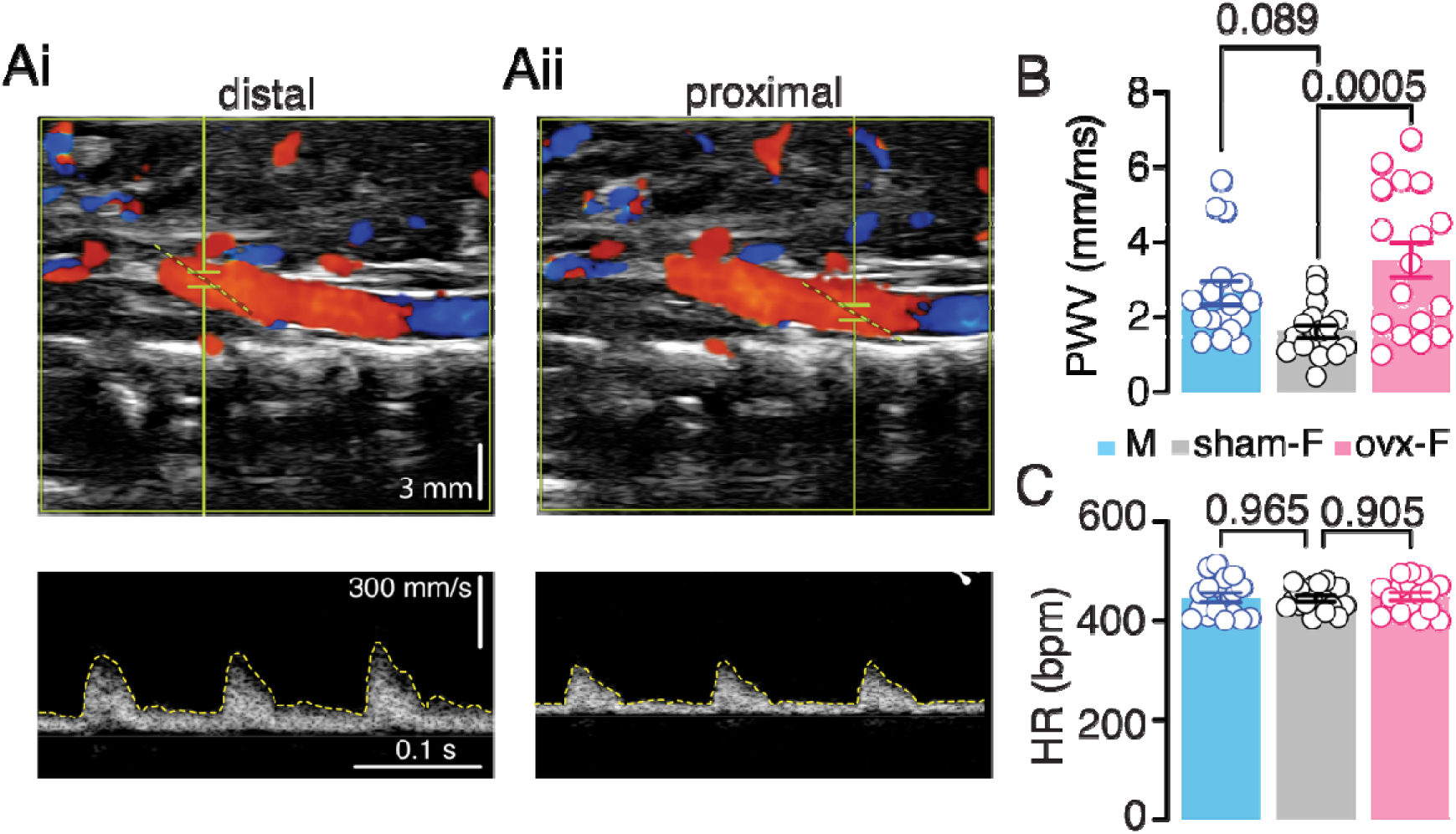
*In vivo* aortic pulse wave velocity (PWV) and Doppler-derived aortic flow parameters from male, sham female, and ovx females. Representative ultrasound/Doppler images of the **A)** distal and **B)** proximal aorta (top) and Doppler velocity waveforms (bottom). Bar plot of calculated **C)** PWV and **D)** heart rate. Sample size: male, 17; sham, 18; and ovx, 17. Data represent mean ± SEM. *P*-values are indicated in the figure. One-way ANOVA with Tukey’s multiple comparisons.

## DISCUSSION

The present study uncovers a previously unrecognized layer of sex-based differences in VSM cAMP signaling: sexually dimorphic subcellular cAMP compartmentalization at the SR. Using targeted FRET biosensors, proximity ligation assays, Ca^2+^ spark imaging, functional myography and PWV, we show that sham females sustain a strong cAMP pool at the SR after β-AR stimulation, while males and ovx females do not. This variation is linked to different PDE3 and PDE4 distributions at the SR, decreased Ca^2+^ spark amplitude, and reduced β-AR vasodilation in males and ovx females. Overall, these results support a model in which PDE activity at the SR is a critical factor in sex-specific regulation of VSM reactivity.

The concept that cAMP does not diffuse freely throughout the cell but is instead organized into spatially discrete pools has transformed our understanding of how this ubiquitous second messenger achieves signaling specificity (Bers et al., 2019; Zaccolo and Kovanich, 2025). In cardiac muscle, compartmentalized cAMP pools, shaped by the strategic localization of PDEs at β-AR targets at different subcellular sites, are well characterized and functionally important for fine-tuning neurohormonal responses in the heart (Bers et al., 2019; Fu et al., 2024; Surdo et al., 2017). In VSM, cAMP compartmentation has been examined mainly at the plasma membrane and in bulk cytosol, whereas the existence and regulation of a functionally distinct SR-associated cAMP pool remain poorly defined (Zhai et al., 2012). Moreover, although sex- and age-dependent differences in microvascular PDE expression have been reported (Wang et al., 2021), and sex-dependent cAMP compartmentation has been suggested in myocardium (Caldwell et al., 2023; Machuki et al., 2019), whether cAMP is spatially organized in a sex-dependent manner in VSM remained unknown. Our data directly address this gap. Using compartment-targeted FRET biosensors (ICUE3 and CUTie), we show that β-AR stimulation produces cAMP accumulation across all subcellular compartments in sham female VSM but is largely excluded from the SR in males. This finding demonstrates that subcellular cAMP compartmentation differs by sex and identifies the SR as a sexually dimorphic cAMP signaling domain in VSM.

Subcellular PDE signalosomes are key determinants of local cAMP gradients and cellular function (Bers et al., 2019; Surdo et al., 2017). Our data show that PDE4 shows a greater association with the SR in male VSM than in sham females, as detected by PLA. In VSM from ovx females, both PDE3 and PDE4 show increased association with the SR. These findings suggest that increased PDE tethering at the SR creates a high-capacity cAMP hydrolysis zone around the RyR, effectively insulating this microdomain from bulk cAMP signals. In sham females, reduced PDE-RyR association would allow cAMP within the SR microdomain, where it can engage PKA and modulate RyR function. This model is consistent with the biosensor data and provides a mechanistic basis for the sex difference in SR cAMP pools. Notably, males and ovx females exhibit distinct PDE distribution patterns. Males exhibit predominantly higher PDE4-RyR association, whereas ovx females show elevations in both PDE3 and PDE4 at the SR. Yet both groups converge on a similar attenuation of the SR cAMP pool and a similar vasodilatory phenotype. This convergence suggests that the functional consequence depends on aggregate PDE activity at the SR microdomain rather than on the specific isoform responsible. Whether PDE3 and PDE4 play redundant or complementary roles at the RyR in VSM, and whether their relative contributions differ under specific physiological conditions, are important questions for future investigation. A comparable idea has emerged from work examining the microtubule network in VSM, in which colchicine- or nocodazole-induced microtubule disruption enhances isoproterenol-mediated relaxation and hyperpolarization of renal and mesenteric arteries (Jepps, 2021; Lindman et al., 2018). Intriguingly, the microtubule stabilizer paclitaxel prevents this effect without altering the cell’s capacity to produce cAMP (Jepps, 2021; Lindman et al., 2018). Together with our findings, these observations support a broader view in which the spatial organization of the signaling machinery, rather than its abundance, sets the gain of β-AR responses in VSM.

A central finding of this study is that ovariectomy abolishes the female-specific SR cAMP domain, mirroring the male pattern in both biosensor responses and PDE-RyR association. These findings implicate ovarian hormones, potentially estrogen, consistent with evidence that estradiol modulates global cAMP in VSM through rapid nongenomic mechanisms and regulates vascular signaling (Farhat et al., 1996; White et al., 1995; Zhu et al., 2002). Our transcript data showed no differences in PDE expression among male, sham, and ovx females, ruling out ovarian hormone-dependent transcriptional regulation of PDE isoform involvement in sex-specific cAMP responses at the SR. Instead, the data point to post-translational regulation of PDE complexes at the SR. Future studies using selective estrogen receptor agonists and antagonists will be needed to dissect the receptor subtypes and downstream mechanisms responsible for the hormone-dependent organization of PDE activity at the SR.

Functionally, reduced cAMP pool at the SR of male and ovx females was concomitant with significantly lower Ca^2+^ spark amplitude in VSM from males and ovx females than in sham females, whereas spark frequency and kinetics are largely unaffected. Ca^2+^ sparks are a critical link between SR signaling and VSM dilation, and cAMP/PKA regulation of this axis is well documented (Nelson et al., 1995; Porter et al., 1998). The observation that mesenteric arteries from sham females dilate more robustly in response to isoproterenol than those from males or ovx females is consistent with a more robust cAMP pool and Ca^2+^ sparks in VSM from sham females. These data suggest a sex-specific organization of the cAMP signaling machinery downstream of receptor activation. Sex-dependent organization of the β-AR relaxation machinery has also been reported at the level of ion-channel accessory subunits. Kcne4 deletion altered mesenteric artery reactivity in male but not female mice, and female arteries showed lower Kcne4 expression (Abbott and Jepps, 2016). Similarly, G_s_-coupled relaxations in rat mesenteric arteries are sexually dimorphic (Baldwin et al., 2022). Our data extend these observations downstream to the subcellular organization of the second messenger itself. In this model, the enhanced vasodilatory response of sham females reflects a subcellular environment in which β-AR-generated cAMP has unimpeded access to the SR microdomain, where it can engage RyR/PKA-dependent pathways that contribute to smooth muscle relaxation. The attenuation of this response following ovariectomy, and its correspondence with changes in PDE-SR association and spark amplitude, suggests that the integrity of the SR cAMP pool is a hormonally regulated determinant of β-AR vasodilatory capacity. Moreover, ovx mice display increased arterial stiffness. Together, these data indicate that ovariectomy promotes a vascular phenotype characterized by increased arterial stiffness and altered vascular reactivity.

Several limitations of the present study merit consideration. First, we performed our biosensor and spark experiments in mouse VSM from the aorta and mesenteric arteries, respectively, and it remains to be determined whether the same sex-specific compartmentalization patterns apply across vascular beds with different physiological roles and sympathetic innervation densities as well as different species. This consideration is supported by evidence that β-AR-driven relaxation engages distinct downstream effectors in different beds (Chadha et al., 2012). Second, while ovariectomy broadly implicates ovarian hormones, our data do not distinguish the specific contributions of estrogen, progesterone, and their respective metabolites to the observed cAMP compartmentalization phenotype. Third, the precise molecular mechanisms by which ovarian hormones regulate PDE-SR association in VSM remain undefined. Finally, although our data are consistent with a model in which SR cAMP signaling modulates RyR function and likely downstream BK_Ca_-mediated vasodilation, direct evidence for this signaling chain in the context of sex differences, for example, through targeted BK_Ca_ inhibition or RyR phosphorylation measurements, will be needed to fully validate the proposed pathway. Moreover, K_V_7.4/K_V_7.5 channels are additional cAMP-and PKA-sensitive mediators of β-AR relaxation in arterial smooth muscle (Chadha et al., 2012; Jepps et al., 2015), and their contribution to the sex differences reported here warrants further examination alongside BK_Ca_.

In summary, this study highlights that biological sex and ovarian hormone status determine subcellular cAMP compartmentalization in VSM, with sham females maintaining a robust SR cAMP pool that is absent in males and lost after ovariectomy. This sexually dimorphic cAMP architecture is shaped by sex-specific distributions of PDE4 at the RyR microdomain and is associated with differences in Ca^2+^ spark amplitude and β-AR vasodilation. These findings reveal a previously unrecognized mechanism by which sex organizes intracellular signaling in VSM, with implications for understanding sex-based differences in vascular physiology and, potentially, in the vascular response to disease and therapeutic intervention.

## AUTHOR CONTRIBUTIONS

EAPS: lead author who acquired, analyzed, and interpreted data for the work, collected all the data and organized it for the repository, and revised it critically for important intellectual content. IV-A, HV, PS, ND-R, MM-AB: acquired, analyzed, and interpreted data for the work, and revised it critically for important intellectual content. YKX: generated and shared resources for the work and revised it critically for important intellectual content. MFN: conceived and designed the work, analyzed and interpreted data for the work, revised it critically for important intellectual content, edited the manuscript and acquired funding for the work. MN-C: lead author, conceived and designed the work, analyzed and interpreted data for the work, drafted and obtained final approval of the manuscript, and acquired funding for the work. All authors have approved the final version of the manuscript and agreed to be accountable for all aspects of the work in ensuring that questions related to the accuracy or integrity of any part of the work are appropriately investigated and resolved. All persons designated as authors qualify for authorship, and all those who qualify for authorship are listed.

## DATA AVAILABILITY

Data supporting the findings shown here are included in the manuscript and can be downloaded from doi: 10.5281/zenodo.22713346.

## GENERATIVE AI STATEMENT

Claude (Anthropic Opus 4.6) and OpenAI (GPT-5.6 Sol) were used during the preparation of this manuscript for the following reasons. First, we used the LLMs for language editing to improve clarity, flow, and formatting consistency. Second, we used OpenAI GPT-5.6 Sol to generate an artistic graphical abstract based on the data in the manuscript. All figures and graphs were generated in GraphPad PRISM. No figure or graphic was produced with the help of LLM tools. We did not use LLMs to acquire data or generate scientific claims. All AI-assisted text was revised and edited by the authors, who accept full responsibility for the accuracy, integrity, and content of the manuscript.

## NONSTANDARD ABBREVIATIONS AND ACRONYMS

AC: adenylyl cyclase
AKAPs: A-kinase anchoring proteins
β-AR: β-adrenergic receptors
Ca^2+^: calcium
cAMP: 3’,5’-cyclic adenosine monophosphate
CUTie: cAMP Universal Tag for imaging experiments
Cyt: cytosol
G_s_PCRs: G protein-coupled receptors
ICUE3: Epac1-camps-based FRET sensor
ISO: isoproterenol
ovx: ovariectomized
PDE: Phosphodiesterase
PKA: protein kinase A
PM: plasma membrane
PWV: pulse wave velocity
SR: sarcoplasmic reticulum
VSM: vascular smooth muscle

